# An In Vivo Model of α-Synuclein Spread from Gut to Brain

**DOI:** 10.1101/2025.03.13.643189

**Authors:** Lisa M. Barnhill, Sataree Khuansuwan, Wael El Nachef, Sung Min Ha, Marisol Arellano, Kazi Md Mahmudul Hasan, Aaron Kim, Jeff M. Bronstein

## Abstract

**Background:** Parkinson’s disease is a progressive neurodegenerative disorder characterized by the presence of pathological aggregation of the protein alpha-synuclein and the loss of dopaminergic neurons in the substantia nigra. There is evidence that misfolding and propagation of alpha-synuclein aggregates through networks of interconnected neurons is responsible for the pathological spread and progressive neuron loss. However, *in vivo* models demonstrating such pathological progression remain elusive.

**Results:** This study utilizes a zebrafish model in order to interrogate the mechanisms of alpha-synuclein toxicity and spread. We describe the development of a zebrafish model of endogenous neuronal human alpha-synuclein expression that causes, in young fish, behavioral and neuronal changes as well as microglia activation. In aged fish, alpha-synuclein expression induces a slow but progressive pathological phenotype manifesting in neuron loss within the gut and the CNS. This model is further utilized to seed gut pathology by incorporating a novel method of feeding human alpha-synuclein preformed fibrils in order to initiate protein misfolding at an early age. The combination of endogenous neuronal expression of alpha-synuclein and the exogenous addition of misfolded protein facilitates the development of brain pathology and subsequent neuron loss in the CNS. In addition to the pathological alterations induced with the fibril feeding model, genetic changes were identified by single cell RNA sequencing. These gene changes resulted in pathway alteration that implicate neurodegenerative disease processes.

**Conclusion:** This model of alpha-synuclein pathology is useful for understanding mechanisms underlying disease initiation and can replicate the progressive development of pathological synuclein accumulation. It has the potential to induce neuron to neuron spread and also offers a way to explore what interventions may prevent such pathological progression.

## Introduction

Most neurodegenerative disorders of aging are characterized pathologically by the accumulation of misfolded proteins^1^. In Parkinson’s disease (PD), α-synuclein (αSyn) is the protein that aggregates and forms insoluble aggregates called Lewy Bodies; toxic species of misfolded αSyn are likely central to the pathogenesis of PD ^2^. αSyn is a relatively abundant protein in neurons and, although its function is not completely understood, it appears to play a role in regulation of synaptic vesicle trafficking and recycling. Normally, αSyn exists as an unfolded monomer, but under certain conditions, it can misfold and form soluble oligomers and insoluble fibrils ^3^. These misfolded versions of the protein can then seed further αSyn misfolding and accumulation, thus propagating abnormal protein species. Furthermore, misfolded aggregates and oligomers can spread to neighboring neurons and perpetuate this pathological process through networks of interconnected neurons. These events, termed templating or spreading, are similar to the formations initiated by prion proteins and appear to underlie most, if not all, common neurodegenerative disorders ^3–5^.

There is evolving evidence that α-Syn spreading occurs in PD. Based on clinical symptoms and pathology, misfolding appears to start in the gastrointestinal (GI) tract or the olfactory bulb and then spread to the brainstem, substantia nigra, and the cerebral cortex ^6^. Constipation and decreased smell are common symptoms patients experience years prior to onset of the cardinal motor symptoms of PD ^7,8^. Importantly, αSyn pathology has been reported in the GI tract and olfactory bulb years before the diagnosis of PD is made, and these prodromal symptoms may reflect αSyn pathology ^6,9–11^. As αSyn pathology spreads to different regions of the CNS, so do PD symptoms. Human data and animal studies also indicate that the gut-brain connection facilitates the spread of pathological αSyn from the periphery to the CNS via the vagal nerve; in fact, severing this connection can even reduce risk of PD, as shown after truncal vagotomy ^12–15^.

A number of mechanisms have been proposed by which αSyn and other aggregated proteins can spread from one cell to another. Cells can release αSyn as free protein by exocytosis, through the formation of exosomes that are subsequently taken up by neighboring cells, and/or through tunneling nanotubes connecting cells ^16,17^. Among these mechanisms, exosomal spread appears to contribute significantly to aggregated protein propagation in neurodegenerative disorders, but this process is very difficult to study *in vivo* ^4,18^. In order to determine the mechanism of αSyn spread *in vivo*, we developed a zebrafish model of human αSyn overexpression in neurons that causes very little baseline neuronal toxicity. When this transgenic linen is combined with a novel method of introducing human preformed αSyn fibrils through ingestion, the model allows for dynamic monitoring of gut to brain pathology and neuronal toxicity over time. By establishing a model of αSyn propagation, it is possible to interrogate and attempt to interrupt this critical early disease process.

## Materials and Methods

### Transgenic lines

The αSyn expressing ZF were generated using constructs made via Gateway cloning protocols using the pDestTol2CG2 destination vector and p5E(4nrUAS), pME(αSyn), and p3E(T2A-eGFPpA) vectors. Wild type ZF embryos (AB strain) were injected within the first 30 minutes of fertilization with 25pg of vector pDestTol2CG2[4nrUAS-αSyn -T2AeGFPpA] along with 25pg of tol2 transposase mRNA. Germline insertion was screened using a cardiac GFP marker and the F1 generation was screened by outcrossing to Tg[HuC:Gal4] in order to confirm brain positive GFP expression from UAS. The Tg[UAS-αSyn -T2A-eGFP] line used in this study has been shown to contain a single insertion through multiple generations and has been confirmed to express both αSyn and eGFP in HuC-positive neurons within the CNS through antibody labeling.

### Zebrafish behavior

Behavioral analysis was conducted on αSyn + and αSyn - zebrafish embryos at 7dpf in 96 well plates and at 4wpf in 24 well plates; data was collected using the Zebrabox observation chamber and behavior tracking system (Viewpoint). Larval and adolescent fish were acclimated to dark conditions for 10 minutes and then video and data recording was conducted for 60 minutes of light cycling with 10 minute lights on followed by 10 minutes lights off. Distance moved was summed for each 10 minute interval and graphed over the 1 hour behavioral collection. Each plate run at 7dpf had between 15-20 larvae per condition and the behavior assay was run 3 times using different clutches of fish for a final number of 50 fish per condition. At 4wpf, 6 fish per condition were tested in a single run.

### Single cell RNA-seq (scRNA-seq)

Homozygous HuC:Gal4 ZF were crossed with heterozygous UAS: αSynT2AeGFP ZF and embryos were screened at 24hpf for GFP expression in the brain. Thirty αSyn – and 30 αSyn+ larvae (5dpf) were deeply anesthetized with tricaine methanesulfonate, the brains were dissected and placed in 1ml cold Ringer’s Solution on ice. They were washed with DPBS and 500ul room-temperature Accumax Cell Dissociation Solution (Innovative Cell Technologies, San Diego, CA) and incubated in a 37 °C. The brains were triturated with a fire-polished glass Pasteur pipette 20 times every 10 minutes for a total of 40 minutes. The tubes were then placed on ice and 500ul of DPBS was added. The suspension was then filtered through a 40um mini cell strainer (pluriSelect, El Cajon, CA) into a sterile, round-bottom, 14mL (17 x 100mm) Falcon collection tube (Becton Dickinson, Franklin Lakes, NJ). The cells were pelleted at 1100 rpm for 10 minutes at room temperature and resuspended in 70ul of 1xPBS + 0.04% BSA. Samples with a viability above 70% were used for the study. 10X Genomics scRNA-seq was performed by the UCLA Technology Center for Genomics and Bioinformatics (TCGB). 10,000 cells were targeted per run with 50,000 reads per cell on the NextSeq 500 High Output, with the read length being 1x75.

### scRNA-seq Analysis

Bioinformatic analysis was performed as described previously^19^. Brierfly, reads generated from the NextSeq 500 High Output were transferred to the UCLA Hoffman2 Cluster. The 10X CellRanger version 3.1.0 software was used to prepare a zebrafish reference genome for alignment, using Danio_rerio.GRCz11.101.chr.gtf and Danio_rerio.GRCz11.dna.primary_assembly.fa, and the sequence for human alpha-synuclein was added. CellRanger count was run for each GEM well, a total of 3 replicates per condition. CellRanger aggr was run to aggregate the count data.

On RStudio Desktop, “Seurat”, “dplyr”, “ggplot2”, and “patchwork” packages were loaded. The aggregated data was analyzed using Seurat 4.0.3 on R version 4.1.0 “Camp Pontanzen” ^20^. The filtered_gene_bc_matrices files from cellranger aggr were loaded to RStudio, and “Read10X” and “CreateSeuratObject” were used to create Seurat objects for each condition. For each object, cells that had less than 500 detected genes were filtered out. “NormalizeData” was used to log normalize the feature expression measurements for each cell, and “FindVariableFeatures” was used to find variable features for each Seurat object. Then, “FindIntegrationAnchors” was used to identify integration anchors using the Seurat objects as input, with the dimensionality of 1:20 as determined by “ElbowPlot”. These integration anchors were passed on to “IntegrateData”, which resulted in a Seurat object that contained the integrated, batch-corrected matrix of all cells for downstream joint analysis. A linear “scaling” transformation was applied using “ScaleData” to a variance of 1 and a mean of 0. We performed principal component analysis on the highly variable genes in the scaled data using “RunPCA”. Further dimensional reduction was performed using top 20 PCs with “FindNeighbors”, and “RunUMAP”. “FindClusters” was run to cluster cells with 0.5 as a resolution value. “DimPlot” was used to visualize clusters “FindConservedMarkers” was used to identify canonical cell type marker genes conserved across conditions for each cluster. The cell identity of each cluster was determined through the identification of 3-5 cluster markers.

Differentially expressed genes were determined by first creating a column in “meta.data” to hold the cell type identification and treatment information, and switching the “current.ident” to that column. Then, “FindMarkers” was used to find the genes that are differentially expressed. For subcluster analysis, the data was scaled, and was followed by “RunPCA,” “RunUMAP”, “FindNeighbors”, and “FindClusters”. Then, “FindConservedMarkers” was used to identify marker genes for each subcluster. Pathway analysis was conducted using Enrichr.

### Zebrafish brain immunohistochemistry and neuron quantification

5-7dpf immunohistochemistry: Larvae were deeply anesthetized in tricaine methanesulfonate before fixing in 4% paraformaldehyde overnight at 4°C rocking and washed in PBS. Tissues were then permeabilized in a solution of 10ug/mL Proteinase K for 5-7 minutes. Tissue was blocked in 5% lamb/5% donkey serum and labeled with primary antibodies against mCherry (1:500 rabbit-anti-mCherry, PA5-34974 Invitrogen), TH (1:250 mouse-anti-TH, MAB318 Chemicon), and GFP (1:500 chicken-anti-GFP, A10262 Invitrogen).

4wpf immunohistochemistry: At feeding study conclusion around 4wpf, deeply anesthetized in tricaine methanesulfonate before fixing in 4% paraformaldehyde overnight at 4°C rocking and washed in PBS. Brains of fixed fish were dissected out and cleared in a modified Clarity solution (0.2M boric acid, 4% SDS, 20mM lithium hydroxide) by incubating tissue in Clarity at room temperature rocking for 4 days followed by washes in PBS. Tissue was then blocked for 4 hours in 5% lamb/5% donkey serum and labeled with primary antibodies against TH (1:250 mouse-anti-TH, MAB318 Chemicon), GFP (1:500 chicken-anti-GFP, A10262 Invitrogen), pSer129 (1:250 rabbit-anti-pSer129, 23706 Cell Signaling Technologies), or αSyn (1:250 mouse-anti-synuclein, 610787 BD Biosciences).

After washing primary antibody, tissue was incubated with secondary antibody overnight at 4°C diluted as follows (1:500 goat-anti-chicken 488, A11039 Invitrogen; 1:500 goat-anti-mouse 568, A11004 Invitrogen; 1:500 goat-anti-rabbit 647, A21244 Invitrogen).

All labeled tissue was cleared in glycerol and imaged on a Leica SPE confocal microscope with a 40x oil objective. TH neuron quantification was conducted on randomized and blinded images for diencephalon and telencephalon counts through the stacks using ImageJ.

### Microglia quantification

ZF were assessed for microglial activation at 5dpf as previously described^19^. Briefly, 5dpf *mpeg1*:mcherry larvae were labeled via immunohistochemistry as described above and were imaged on a Leica SPE confocal microscope with a 40x oil objective. Images were projected and converted to greyscale using ImageJ. Binary images were then skeletonized and activation was quantified by measuring branch length using Analyze Skeleton and Simple Neurite Tracer.

### Zebrafish gut immunohistochemistry and neuron quantification

Adult ZF between 1-2.75 years of age were labeled and imaged for gut neuron quantification as described previously ^21^. Briefly, fish were euthanized before gut tissue was dissected and imaged to determine length using ImageJ software. A 10uL pipette tip was ensheathed in the intestine to maintain a straight configuration and then placed in 4% paraformaldehyde overnight at 4°C to fix. After removal from the pipette tip, tissue was incubated in blocking solution (2% goat serum, 1% bovine serum albumin, 1% DMSO, 0.1% TritonX 100, and 0.05% Tween) followed by primary and secondary antibody labeling (1:200 anti-HuC/D, A21271Thermo Fisher Scientific; 1:500 goat-anti-mouse 647, A21242 Thermo Fisher Scientific). Intestinal tissue was optically cleared in refractive index matching solution (RIMS) for 48 hours and mounted in low-melt agarose (IBI Scientific, IB70056) for imaging on Zeiss Z.1 Light dual sided illumination sheet fluorescence microscope using ZEN software. Images were tiled followed by manual stitching and automated cell counting with the “Spots” feature on Imaris x64 v9.6.0 (Bitplane).

### Zebrafish protein extraction and Western blot

Adult ZF were deeply anesthetized in tricaine, decapitated, and brain tissue was manually dissected with forceps. Tissue was frozen at -80°C before homogenization in 1xRIPA with protease inhibitors. Protein preps were briefly sonicated on ice before centrifugation at 15,000rpm and supernatant was stored at -80C. 40ug of protein, as determined by BCA assay, was run on a 12% Bolt Bis Tris gel for 50min at 150volts, transferred to a nitrocellulose membrane, and blocked in 5% non-fat milk. Primary antibodies (1:800 rabbit-α-Elavl3+4, GTX128365 GeneTex; 1:2500 mouse-α-tubulin, T9026 Sigma Aldrich) as well as secondary antibodies (1:2500 mouse-anti-rabbit HRP, sc-2357 Santa Cruz Biotech; 1:2500 goat-anti-mouse HRP, 62-6520 Invitrogen) were diluted in 1% non-fat milk. Blots were developed in ECL Plus substrate for 5 minutes before imaging on Azure Biosystems imager and band intensity was quantified on ImageJ using the Analyze Gels tool.

### Human αSyn preformed fibril-enriched zebrafish food

Modified diets were created for ZF between the ages of 5dpf and 4 weeks post fertilization (wpf). Human alpha-synuclein preformed fibrils (αSyn PFF) were prepared as previously described^22^. Fibrils were stored at -80°C until use; they were diluted in water to a concentration of 250ug/150uL before being sonicated on ice once per second for 1 minute using a probe sonicator. Gemma Micro 75 feed (Skretting Zebrafish) was weighed and aliquoted into 0.3g of feed per feeding experiment. 150uL of either water or sonicated fibril mixture containing 250ug PFF were added to the 0.3g of powdered food and mixed to completely wet the feed. The paste was then spread in a sterile dish and dried overnight. Dried powder was then pulverized until similar in size to the original feed, placed in microfuge tubes with approximately 0.1g/tube, and frozen at -80°C with desiccant until use.

Feeding experiments began with equal numbers of larvae in 200mL of E2 water at 5dpf (18-30 larvae per tank). These larvae were fed small amounts of enriched food once per day until 4wpf. 0.1g of feed was sufficient to feed 2 tanks for 1 week. Water was cleaned with 50% replacement 2-3 times per week until fixation.

## Results

### αSyn + Transgenic Line Expression Characterization

A novel human alpha-synuclein zebrafish transgenic line was generated through the stable insertion of the construct described in figure 1A (UAS:αSynT2AeGFP). This line can be induced to express αSyn within neurons through crossing to the Gal4 expressing line (HuC:Gal4). Neurons expressing αSyn as identified by antibody labeling experiments generally colocalize with the production of eGFP and exhibit higher expression levels in the forebrain (Figure 1B-E). Fish expressing human αSyn do not exhibit obvious survival or morphological deficits through adulthood, but early behavioral differences in response to light stimulus can be seen in larvae at 7dpf. In general, wild type animals move further when the light is off and reduce their movements when the light is on, thought to be a readout of anxiety^23^. Using an assay measuring light stimulated behavior response, ZF larvae expressing αSyn exhibited reduced distance traveled in dark conditions, but no change in light conditions (Figure 1F-G).

**Figure 1:**
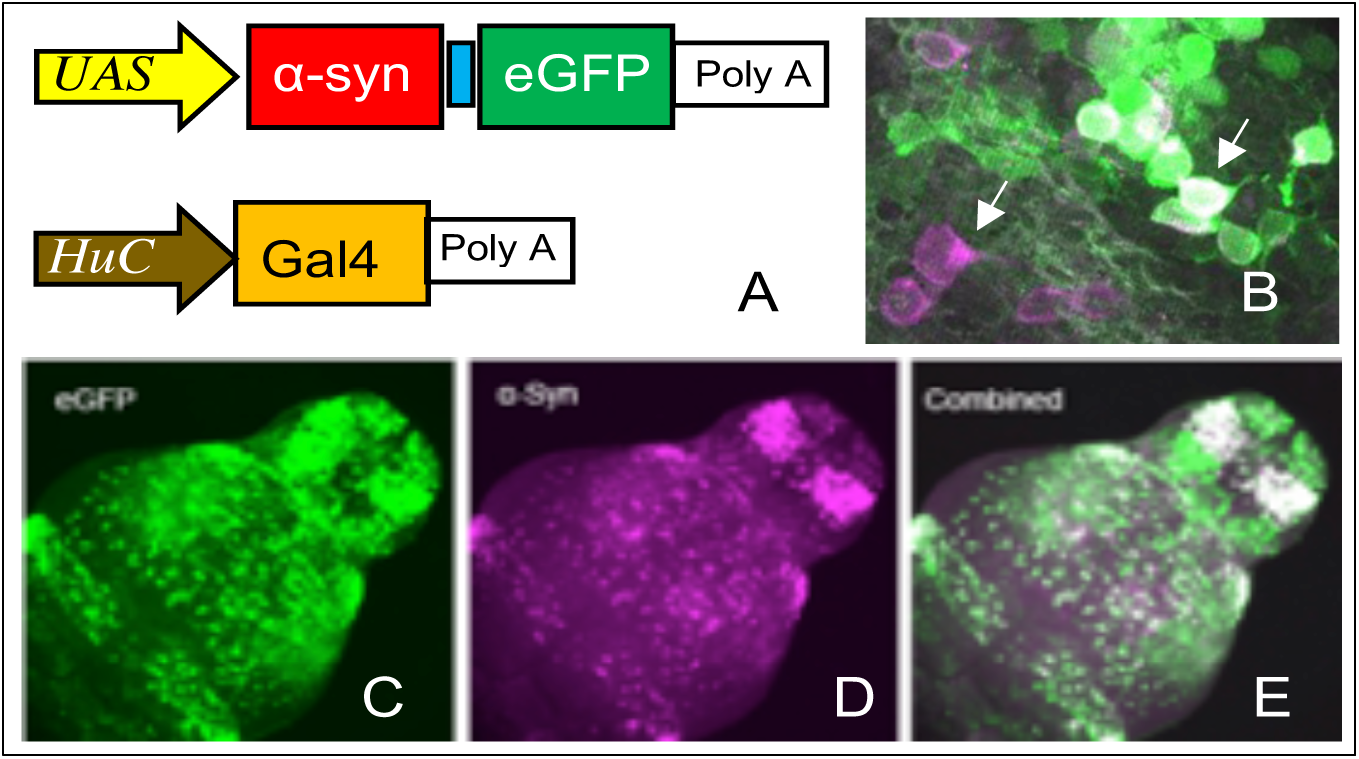
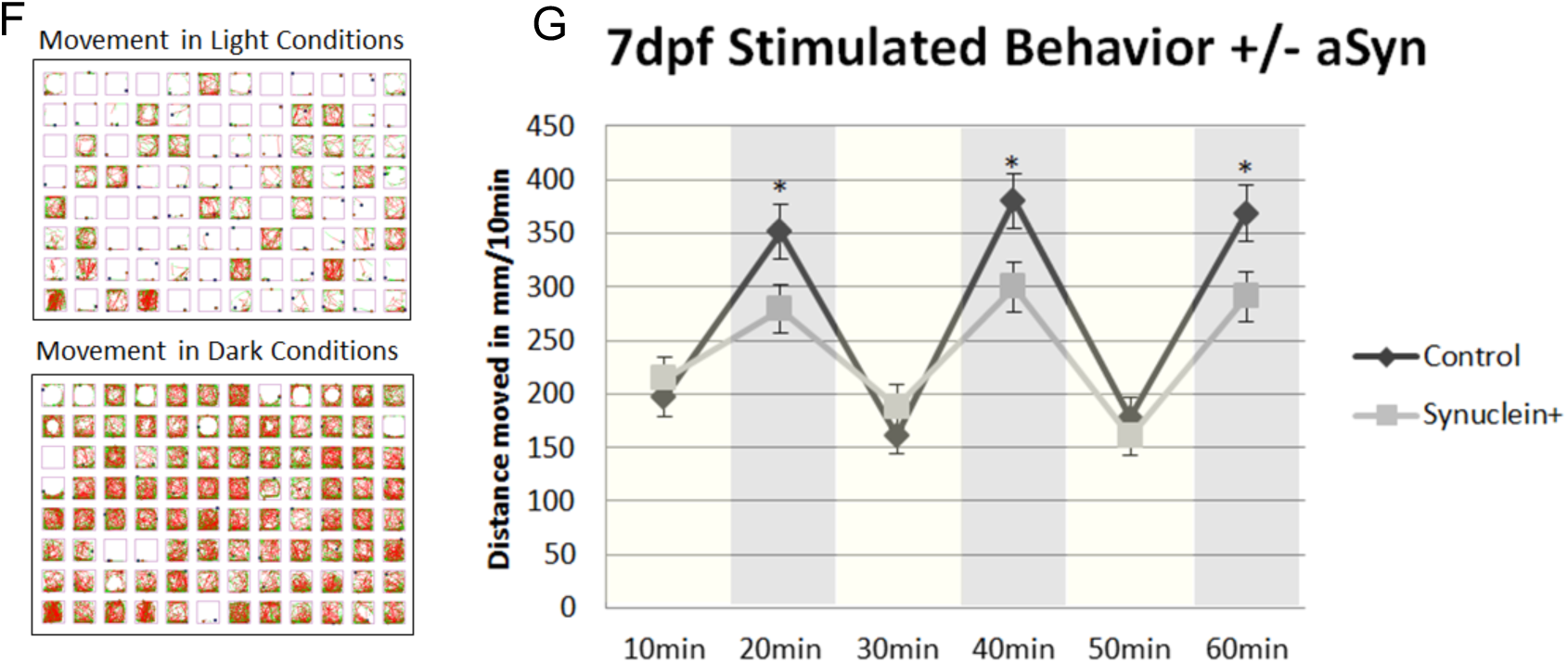
Schematic of the 2 constructs used to generate Tg(HuC:Gal4) and Tg(UAS:αSynT2AeGFP) transgenic lines (A). Neuron expressing eGFP often colocalize with αSyn expression in the 3dpf zebrafish brain (B). Whole-brain expression of eGFP and aSyn are largely colocalized and expressed widely throughout the tissue with areas of significant intensity seen in the telencephalon (C-E). Schematic 96-well plate setup run using the Viewpoint zebrabox (F). Distance moved per 10 minute period across 60 minutes of light dark cycling where yellow columns represent periods of light and grey columns represent periods of dark. Data represents 50 larvae per condition across 3 experimental replicates with 15-20 larvae per run (G).

### Expression analysis comparing αSyn - and αSyn + zebrafish brains

In order to determine how expression of αSyn impacts neuronal expression patterns during development, scRNAseq analysis was conducted on brain tissue from sibling pairs of αSyn – and αSyn + larvae. Three biological replicates resulted in the analysis of a total of 18,916 cells from the αSyn- and 15,194 cells from the αSyn+ larvae. Twenty clusters were identified (Figure 2A) and the genes markers characterizing each cluster are listed in Figure 2B. The 2 largest clusters were of neuronal origin. Neurons were furthered divided into 11 subclusters (Figure 2C) and of these subclusters, cluster 0 had the highest proportion of αSyn expressing cells (Figure 2C). This was the largest subcluster and was enriched with GABAergic neurons (Table 1). Analysis of differentially expressed genes (DEGs) in neuron subcluster 0 from αSyn+ compared to αSyn-fish revealed downregulation of mitochondrial pathways including fatty acid cycling and mitochondrial uncoupling as well as axon subcellular targeting and synapse formation (e.g. reduction in neurofascin, NrCAM and CHL1 interactions) as shown in table 1. The only gene significantly increased in αSyn+ subcluster 0 neurons was mt-atp6, which is essential for mitochondrial function (Table 1). Neuronal subclusters 1 also expressed some αSyn and was composed of predominantly neuronal and retinal progenitor cells. In these neurons, there was a trend for reduced expression of mitochondrial Bcl-2 family and glucocorticoid receptor pathway proteins. Cytoplasmic ribosomal and mRNA processing proteins were increased in this subcluster. Only minor or no significant changes in gene expression were found in clusters 4-10 (Table 1).

**Figure 2:**
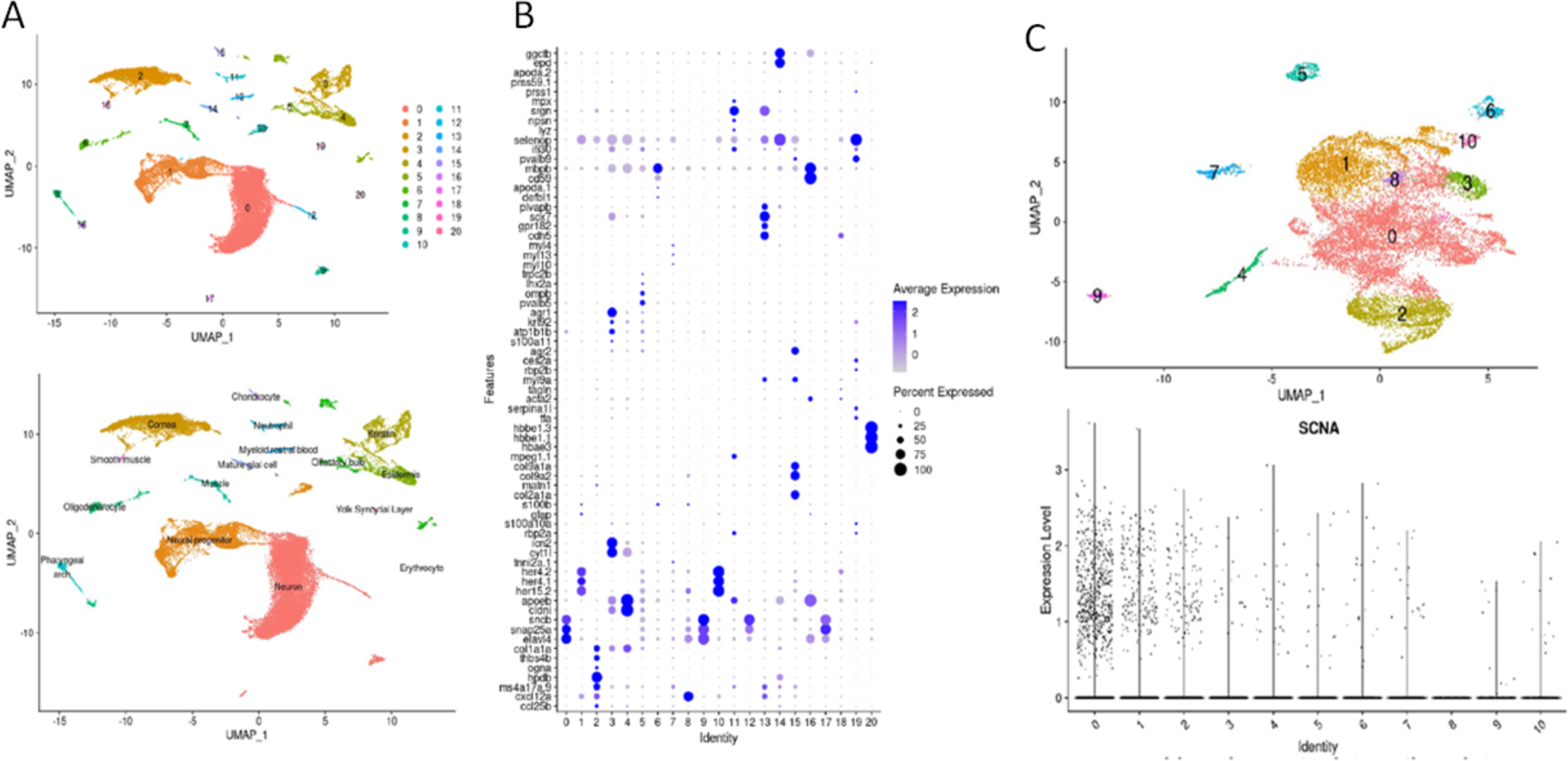
Cluster analysis. A total of 21 clusters were identified and characterized using canonical genes (A and B). A total of 11 neuronal subclusters were identified (C). Of the 11 subclusters, all but cluster 8 expressed appreciable αSyn, with the greatest number of αSyn expressing cells found in cluster 0 (C.)

**Table 1.**
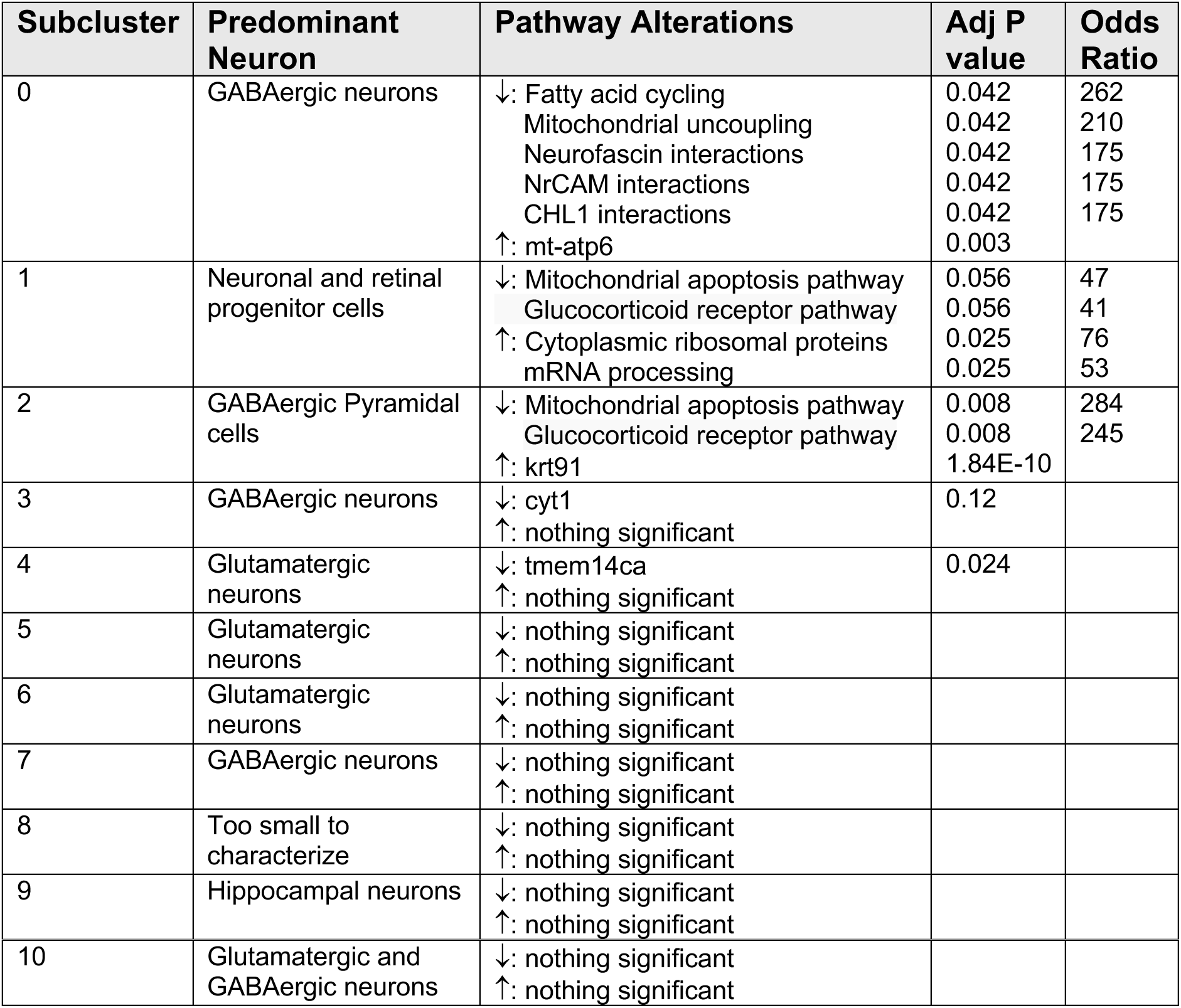

| Subcluster | Predominant Neuron | Pathway Alterations | Adj P value | Odds Ratio |
| --- | --- | --- | --- | --- |
| 0 | GABAergic neurons | ↓: Fatty acid cycling<br>Mitochondrial uncoupling<br>Neurofascin interactions<br>NrCAM interactions<br>CHL1 interactions<br>↑: mt-atp6 | 0.042<br>0.042<br>0.042<br>0.042<br>0.042<br>0.003 | 262<br>210<br>175<br>175<br>175 |
| 1 | Neuronal and retinal progenitor cells | ↓: Mitochondrial apoptosis pathway<br>Glucocorticoid receptor pathway<br>↑: Cytoplasmic ribosomal proteins<br>mRNA processing | 0.056<br>0.056<br>0.025<br>0.025 | 47<br>41<br>76<br>53 |
| 2 | GABAergic Pyramidal cells | ↓: Mitochondrial apoptosis pathway<br>Glucocorticoid receptor pathway<br>↑: krt91 | 0.008<br>0.008<br>1.84E-10 | 284<br>245 |
| 3 | GABAergic neurons | ↓: cyt1<br>↑: nothing significant | 0.12 |  |
| 4 | Glutamatergic neurons | ↓: tmem14ca<br>↑: nothing significant | 0.024 |  |
| 5 | Glutamatergic neurons | ↓: nothing significant<br>↑: nothing significant |  |  |
| 6 | Glutamatergic neurons | ↓: nothing significant<br>↑: nothing significant |  |  |
| 7 | GABAergic neurons | ↓: nothing significant<br>↑: nothing significant |  |  |
| 8 | Too small to characterize | ↓: nothing significant<br>↑: nothing significant |  |  |
| 9 | Hippocampal neurons | ↓: nothing significant<br>↑: nothing significant |  |  |
| 10 | Glutamatergic and GABAergic neurons | ↓: nothing significant<br>↑: nothing significant |  |  |

### Microglial activation after αSyn expression

Brain inflammation and microglial activation is a pathological feature of many neurodegenerative diseases including PD^24^. The transgenic ZF line *mpeg1*:mcherry was used to understand the impact of αSyn expression on neuroinflammation. Microglia that express mcherry were quantified with and without αSyn; in αSyn expressing brains, microglia were distributed differently than in αSyn – siblings (Figure 3A,B,E). There were more microglia located in the telencephalon of the brain, which is the region where αSyn was most highly expressed (Figure 3E). The microglia in αSyn expressing ZF brains had shorter and fewer branches, which is consistent with an activated morphology, as has been previously shown^19^ (Figure 3F). Some of the microglia were in close proximity to αSyn expressing neurons and appeared qualitatively to be engulfing them (Figure 3C-D). A time-lapse video of αSyn expressing fish also expressing mCherry driven by the mpeg1 promoter reveal some microglia that appear to be removing αSyn+ neurites (Supplemental movie 1). These data, when taken together, suggest that ZF expressing human αSyn exhibit a more reactive morphology and inflammatory phenotype, consistent with the conclusion that microglia take on a more activated profile in the presence of αSyn.

**Figure 3:**
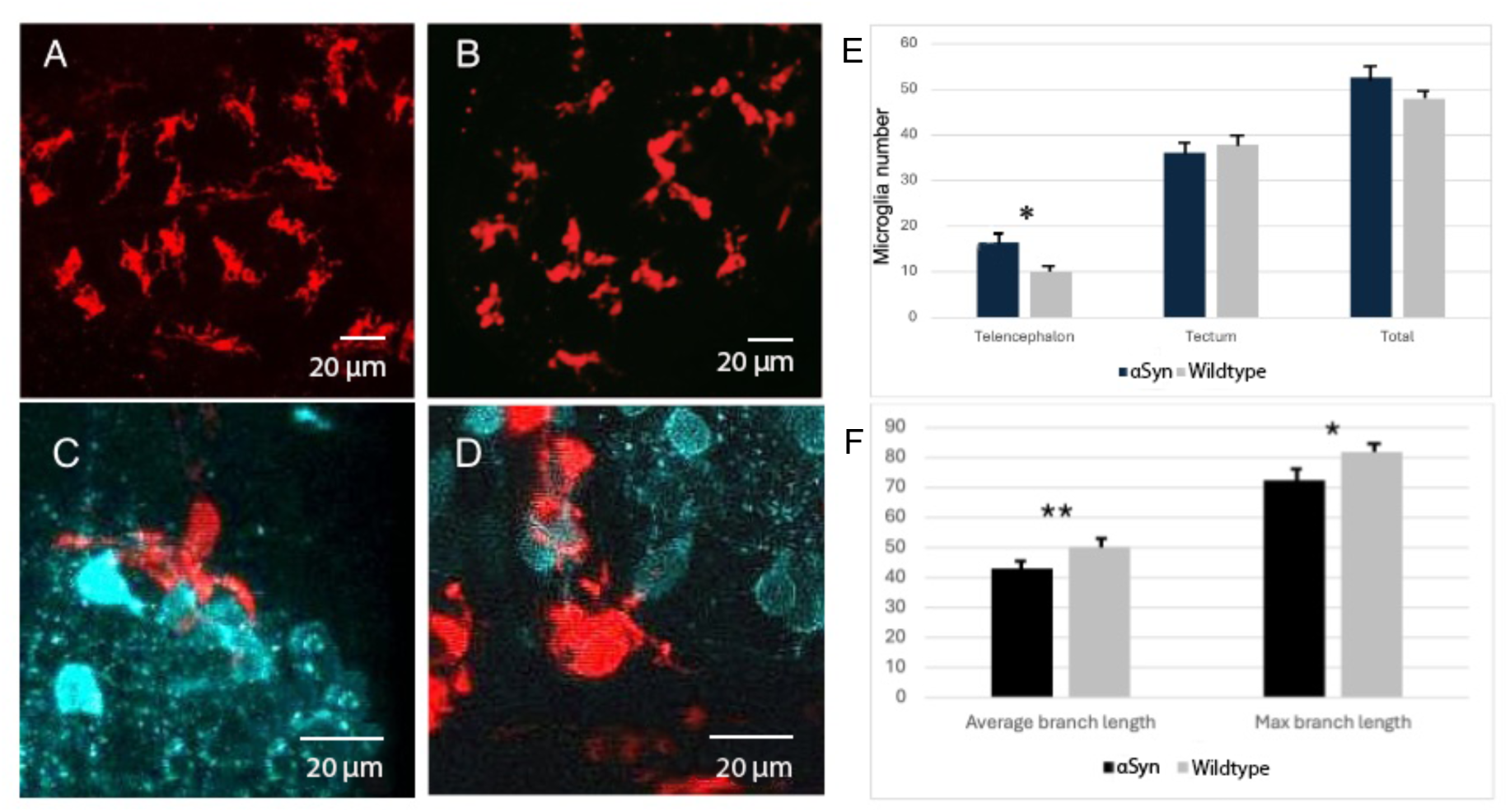
Microglia activation in αSyn expressing fish brains. Microglia αSyn expressing brains (B) were more concentrated in the telencephalon (right) and had longer branches compared to those in wildtype fish (A). Some microglia appear to be consuming αSyn neurons (teal; C and D). N=203 cells from 15 fish for the αSyn and 168 cells from 12 wildtype fish. *= 0.0001 for microglial number, 0.028 for maximum branch length. **=0.009.

### TH neuron loss after aSyn expression

At 7dpf, larvae expressing αSyn in the brain exhibit altered light-stimulated behavior (Figure 1G) as well as inflammatory microglia (Figure 3F). In order to determine if there is any neuronal loss that may accompany these changes, tyrosine hydroxylase (TH)-positive neuronal labeling was conducted in order to quantify dopaminergic neuron number in 7dpf larvae. In larvae expressing αSyn, there was a reduction in the number of TH-positive neurons in the telencephalic region, which contains a high level of αSyn, but no change in the diencephalic region, which has much less αSyn expression (Figure 4A-C).

**Figure 4:**
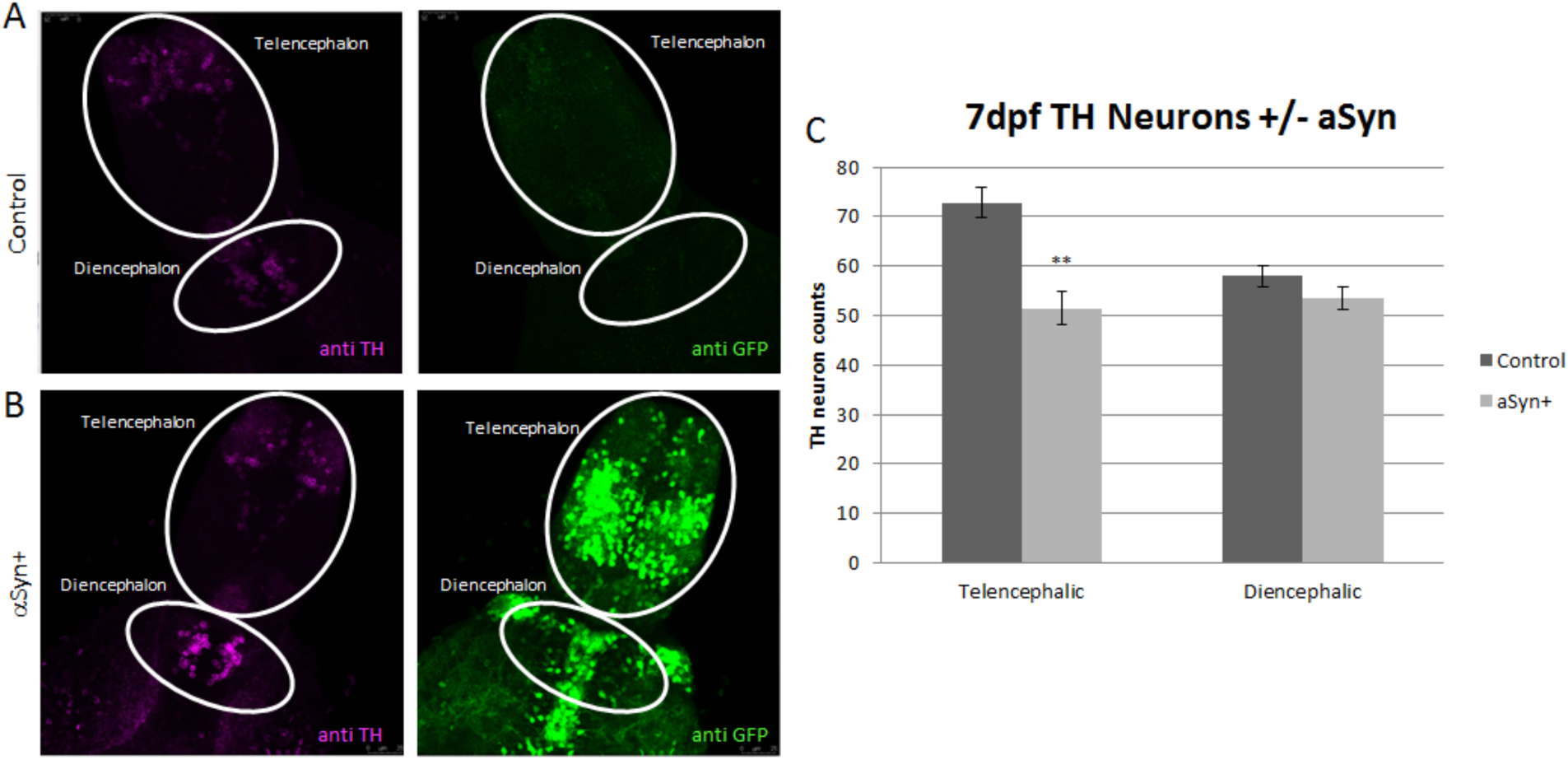
αSyn - (A) and αSyn + (B) larvae were antibody labeled for TH (pink) and GFP (green). At 7dpf, most of the αSyn expression is concentrated in the telencephalon (B). TH neurons in the diencephalic region at this timepoint are unaffected while TH positive neurons in the telencephalon exhibit an approximately 20% reduction in αSyn-expressing fish (C).

### αSyn expression in aged ZF

Although αSyn expressing ZF do not exhibit altered survival or have any overtly toxic phenotype during aging (data not shown), it is important to identify disturbances that may be occurring with chronic overexpression of αSyn that could mirror processes that occur in the early stages of PD. To assess this more carefully, adult αSyn + and αSyn - fish between 6 months and 2.75 years of age were analyzed for HuC/D abundance and gut neuron number to determine the impact of long-term expression. In middle age, around 6 months to 1 year old, the adult ZF expressing αSyn had no changes in enteric HuC/D neuron number (Figure 5A,B) or in brain HuC/D protein expression (Figure 5C,D). However, as the fish aged to between 1.5-2.75 years old, there was a significant reduction in the number of enteric neurons in αSyn expressing fish (Figure 5A,B) as well as a reduction in the amount of HuC/D protein expressed in the brain (Figure 5C,D). The significance of demonstrating progressive neuronal loss is that it more closely recapitulates the process of disease in humans, where overexpression of αSyn results in a slow and progressive degenerative phenotype that occurs both in the gut as well as the brain.

**Figure 5:**
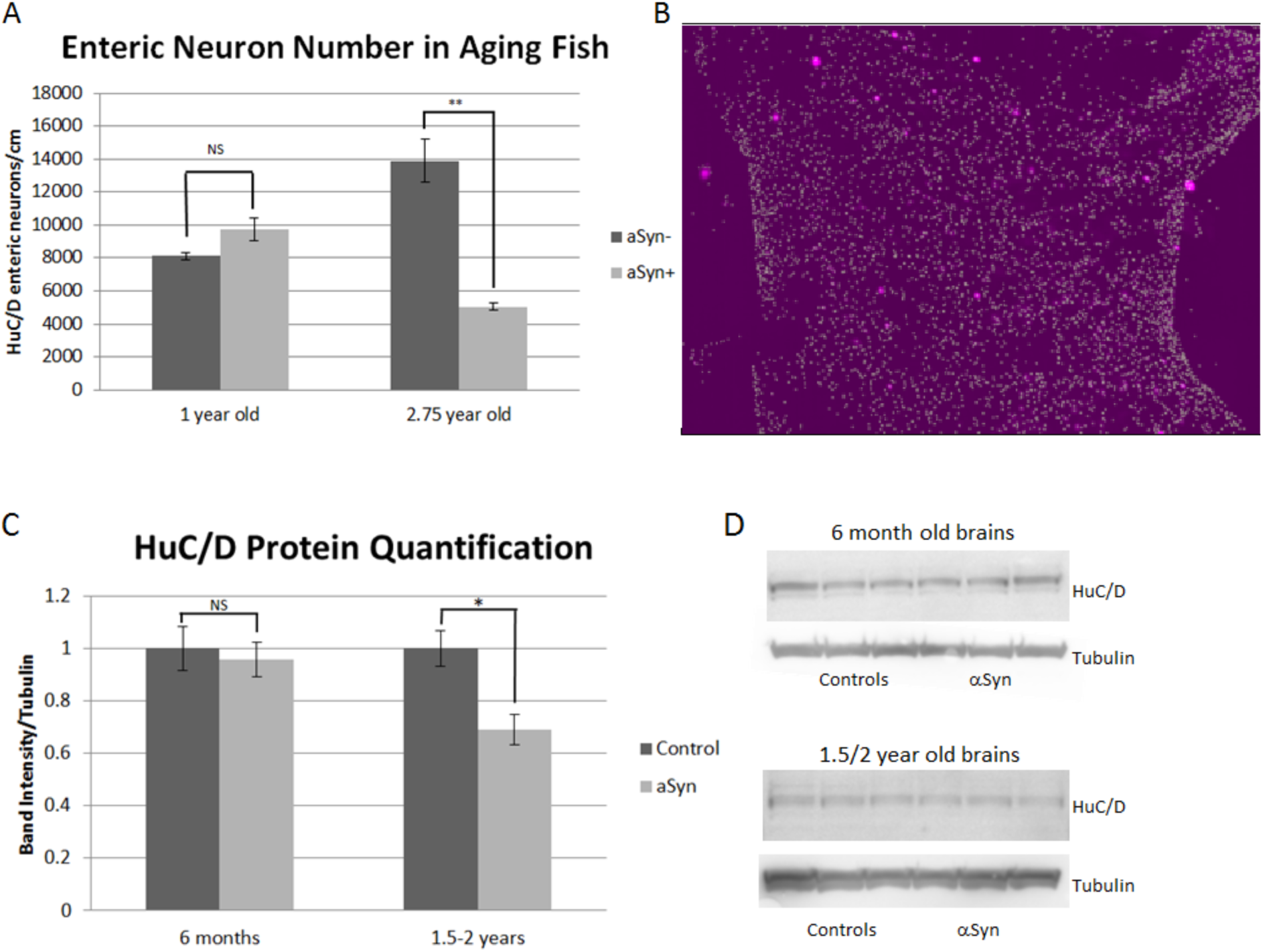
Adult αSyn - and αSyn + fish were aged and labeled for gut HuC/D-expressing neurons (B). Quantification of HuC/D showed a progressive loss of enteric neurons that was evident in aged fish around 2.75 years old (A). Western blot assessment of HuC/D expression in the brains of αSyn - and αSyn + adults demonstrated a reduction in HuC/D protein only in aged fish between 1.5-2 years of age (C,D).

### Model of αSyn fibril feeding in developing ZF

Although alteration in early larval TH neuron number and stimulated behavior were noted in this model, overall toxicity due to αSyn expression is quite minor. There was no overt survival difference between αSyn + and αSyn - fish through at least 1 year of age (data not shown). Additionally, adult behavior, although highly variable, did not exhibit baseline differences between these groups. Because these fish seem quite resilient to αSyn expression, we attempted to challenge the model further by adding human αSyn preformed fibrils (PFF) in order to potentiate misfolding and proteinopathy. To accomplish this, sonicated αSyn PFF were added to the daily food given to larvae starting at 5dpf through 4 weeks post fertilization (wpf) (Figure 6A). Survival on a diet of αSyn PFF-enriched food was unaffected in these experiments and after 4 weeks of a PFF enriched diet, fish exhibited no change in light stimulated behavior compared to non-αSyn expressing siblings on a control diet (Figure 6B-C).

**Figure 6:**
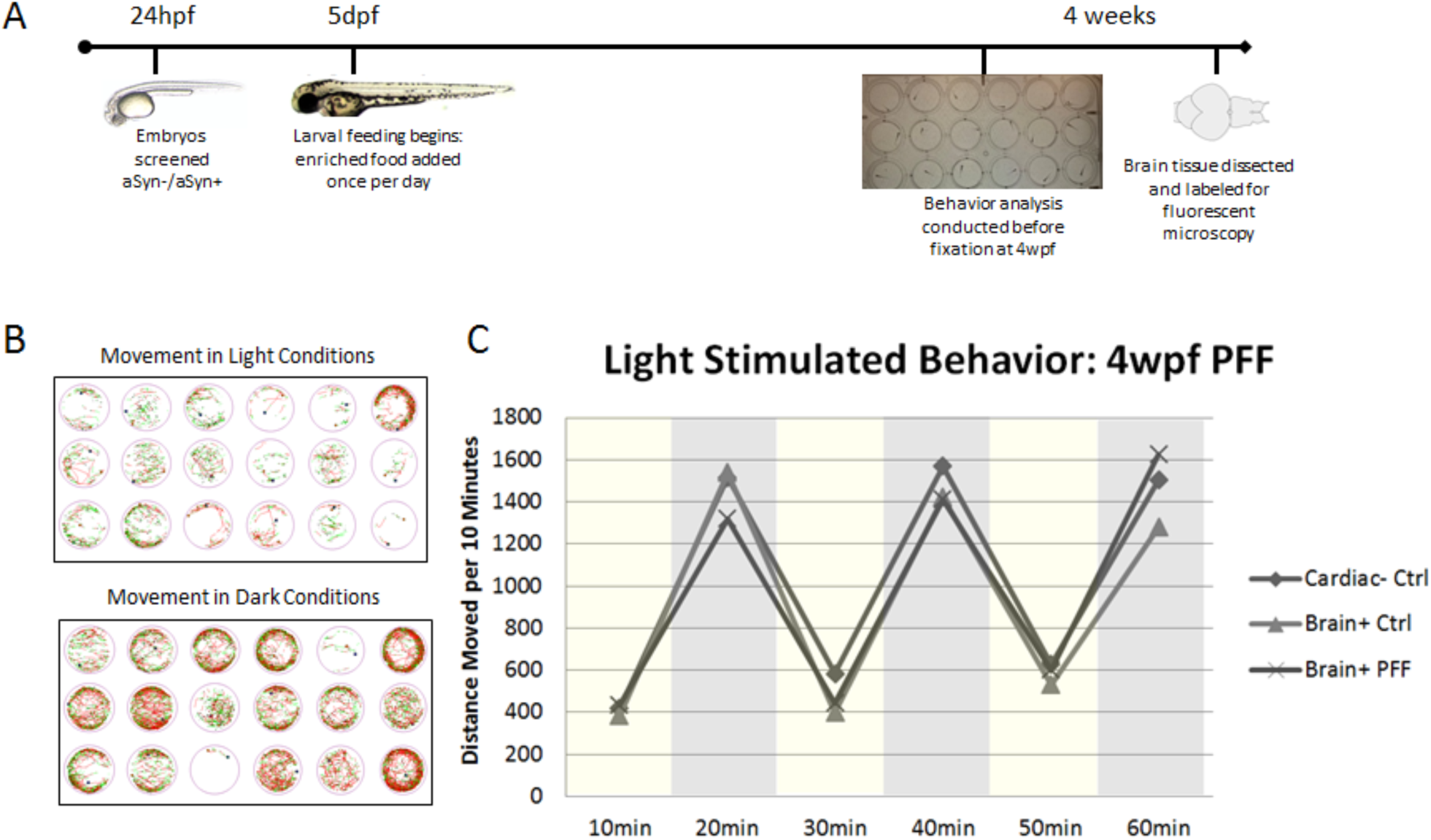
For the fibril feeding experiment, embryos were sorted into αSyn -/αSyn + groups at 24hpf. Starting at 5dpf, tanks of 30 larvae were fed a controlled diet of either powdered food reconstituted in water (control) or powdered food enriched with human αSyn fibrils in water (PFF). Fish were fed once per day until 4 weeks of age (A). Light stimulated behavior was run, and fish were fixed for immunohistochemistry. 4wpf light stimulated behavior was run in 24 wells plates with 6 fish per condition (B). Total distance moved in light vs dark conditions were compared across groups and no differences were seen between αSyn - vs αSyn + or control fed vs fibril fed (C).

### Increased pSer129 labeling and TH neurotoxicity in PFF fed ZF

In order to determine if PFF feeding in αSyn expressing fish caused any change in αSyn accumulation in the CNS, immunohistochemical analysis of pSer129, the phosphorylated αSyn often found in pathological aggregates, was quantified in the brain of PFF fed vs control fed fish^25^. At 4wpf, αSyn ZF^D^fed PFF showed a reduction in TH-positive neurons in both the telencephalon and diencephalon when compared to non-expressing siblings (Figure 7A,B). This further loss of dopaminergic neurons with the addition of PFF shows that the model exacerbates any baseline toxicity that occurs with αSyn neuronal expression. At 4wpf, the minority of GFP-expressing neurons also contain detectable levels of pSer129. After feeding αSyn PFF between 5dpf-4wpf, the percent of GFP neurons also positive for pSer129 more than doubled; ingestion of aggregated αSyn PFF fragments altered the propensity of brain αSyn to become phosphorylated (Figure 7C,D). When taken together, this information shows that ZF consuming a PFF-enriched diet accumulate misfolded αSyn at a higher rate than those on a control diet and additionally, PFF fed ZF show an increase in neuron loss when compared to control-fed siblings. This is likely the result of the spread of misfolded protein from the gut to the CNS and represents a novel *in vivo* model of assessing pathological mistemplating.

**Figure 7:**
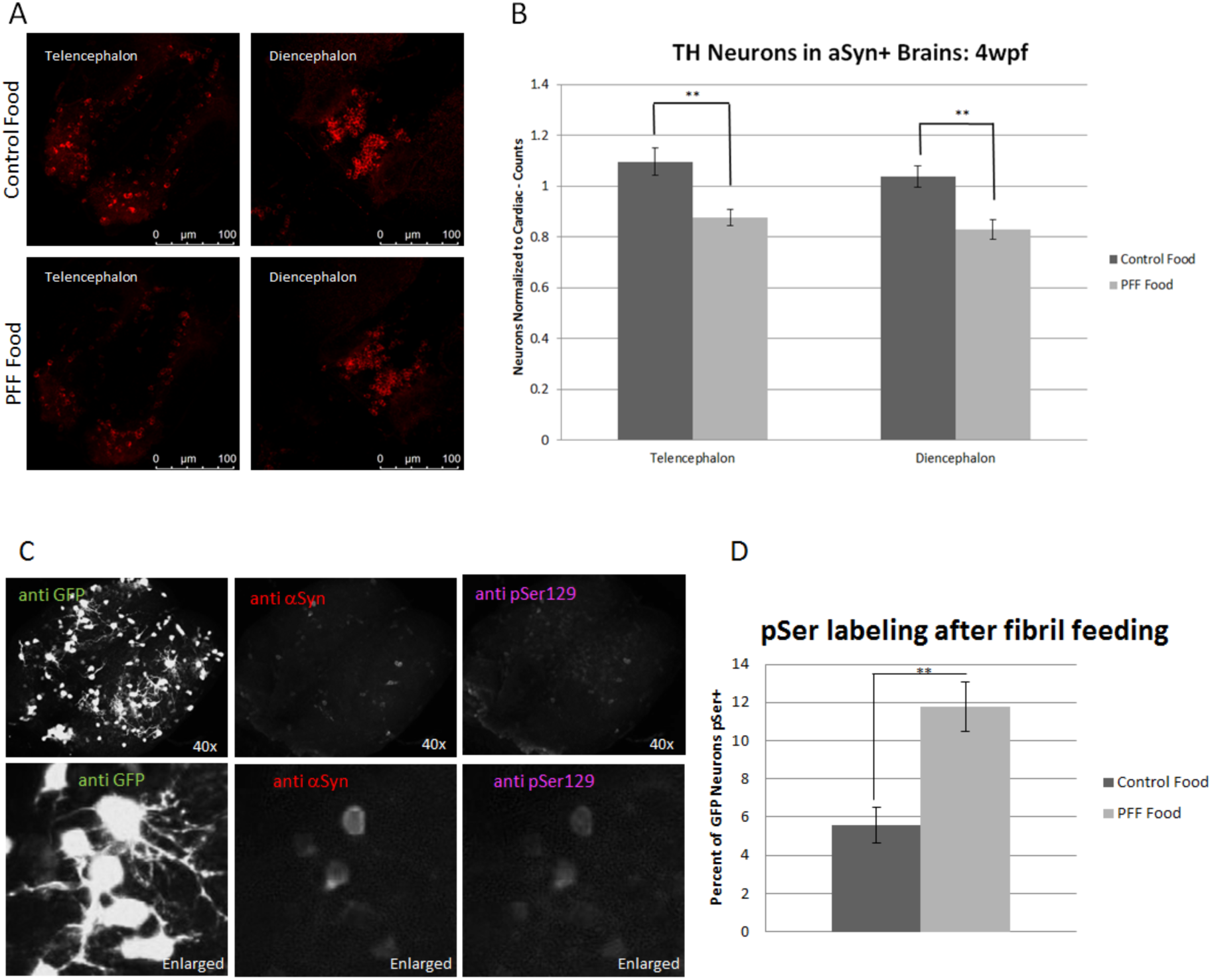
TH neurons were labeled and quantified in the brains of αSyn fish fed either fibril enriched or control food (A). TH neurons in the telencephalon and diencephalon of fish fed PFF enriched food were reduced when normalized to non-αSyn expressing siblings fed the same diet (B). αSyn + brains were labeled for GFP, αSyn, and pSer129 to determine the percent of GFP expressing HuC/D neurons that also contained phosphorylated synuclein after PFF feeding (C). Quantification of pSer129 positive neurons that express GFP showed an increase in pSer129 co-labeling compared to control fed αSyn + siblings (D).

### Transcriptomic analysis of αSyn ZF Fed PFFs

Feeding αSyn-expressing ZF PFFs for several weeks led to a significant increase in pSer129 αSyn presumably by trans-neuronal spread of misfolded αSyn and clearly had an impact on brain. In order to understand the molecular processes that may be underlying such changes, we performed scRNAseq analysis on brains from αSyn-expressing ZF fed a diet enriched with PFF and compared the gene expression to αSyn-expressing ZF fed standard food. The clustering analysis from these brains resulted in 22 total clusters, with each cluster containing cells from both control and PFF fed ZF (Figure 8A,B), which were identified and characterized by the expression of several canonical genes (Figure 8C). Neuronal subcluster analysis on the fibril-fed vs control-fed ZF resulted in 9 neuronal subclusters which were also further characterized by the expression of canonical genes (Figure 8D,E). Among these subclusters, clusters 1, 5, and 8 showed higher levels of αSyn expression (Figure 8F).

**Figure 8:**
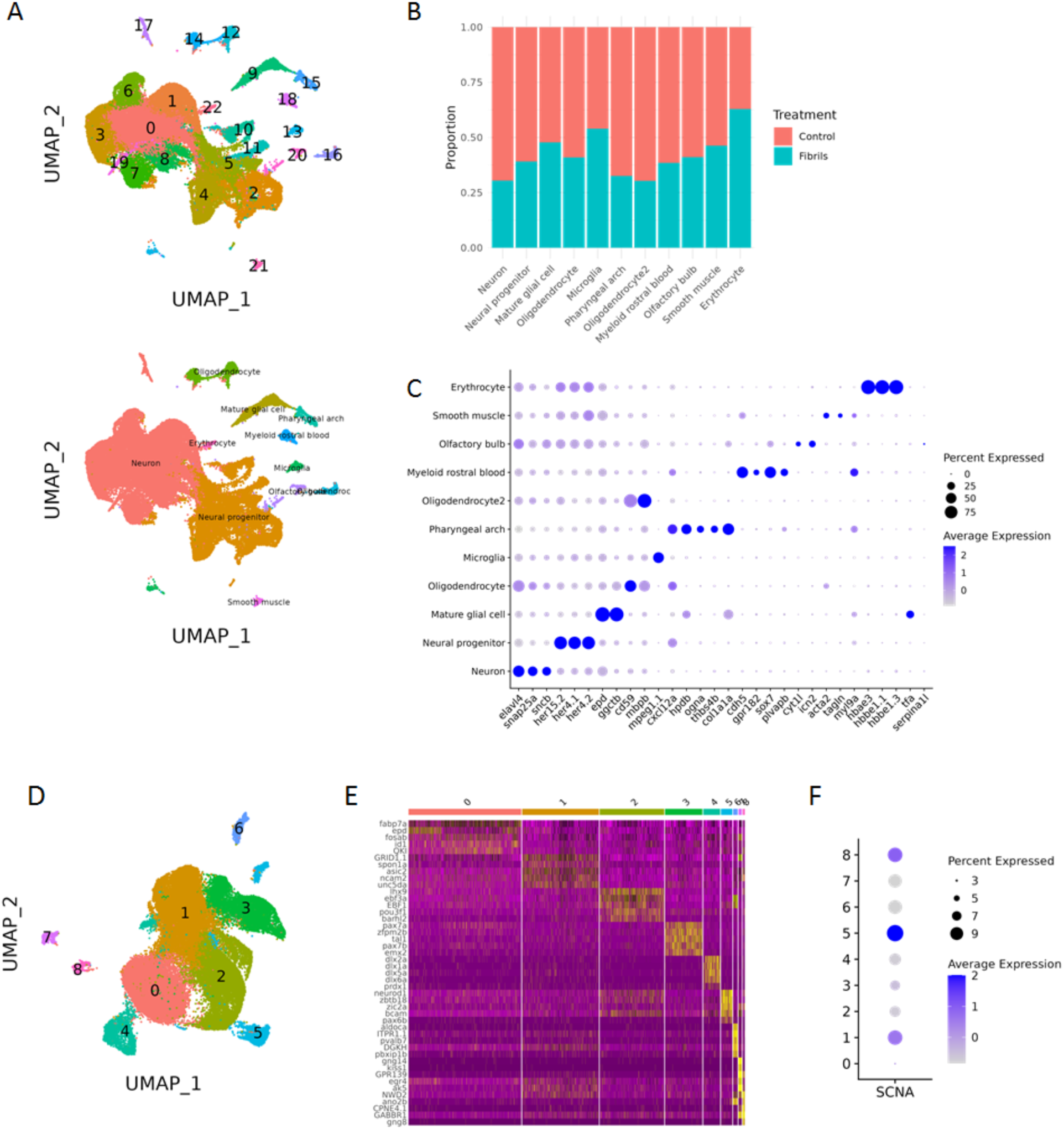
scRNAseq data from the brains of αSyn expressing fish fed a control diet or a diet enriched with PFFs resulted in 22 clusters (A). These clustered were identified and characterized by the presence of canonical genes (C). The proportion of cells representing fibril fed brains were relatively consistent across clusters (B). The neuronal subclustering resulted in 9 clusters (D). Each of these clusters was characterized by the expression of canonical genes (E.) and within the 9 clusters, clusters 1, 5, and 8 had the highest levels of αSyn (F).

Differential gene expression analysis was utilized to identify significant alterations in pathways. Intriguingly, PD- and neurodegeneration-relevant pathways were highlighted in several neuronal subclusters. Pathways-specific changes that were both up- and down- regulated in the main neuronal subclusters after fibril feeding implicated diseases where prion-like spread have been implicated. Coronavirus disease, ribosome, neurodegeneration, Parkinson’s disease, Huntington’s disease, prion disease, Alzheimer’s disease, and ALS, among other pathways, were upregulated in PFF-fed animals in neuron subcluster 0 (Figure 9A). Downregulated pathways in neuron subcluster 1involved neudoegeneration, prion disease, ALS, Huntington’s disease, Parkinson’s disease, and Alzheimer’s disease, among others (Figure 9B). The downregulated pathways in neuron subcluster 2 involved in prion disease, neurodegeneration, Parkinson’s disease, coronavirus, ribosome, ALS,and Huntinton’s disease, among others (Figure 9C). Other clusters showed small changes, primarily involving coronavirus disease and the ribosome (data not shown). These data suggest important changes in the brains of animals fed αSyn fibrils that implicate pathways characteristic of neurodgenerative disease. Many of the differentially expressed genes in these pathways involve mitochondrial oxidative phosphorylation, highlighting that as a potential pathway of interest. It is not clear from these findings whether the pathway changes observed implicate neurotoxic or neuroprotective endpoints, but it is provocative that these neuronal clusters are showing involvement of pathways that may be altered in disease.

**Figure 9:**
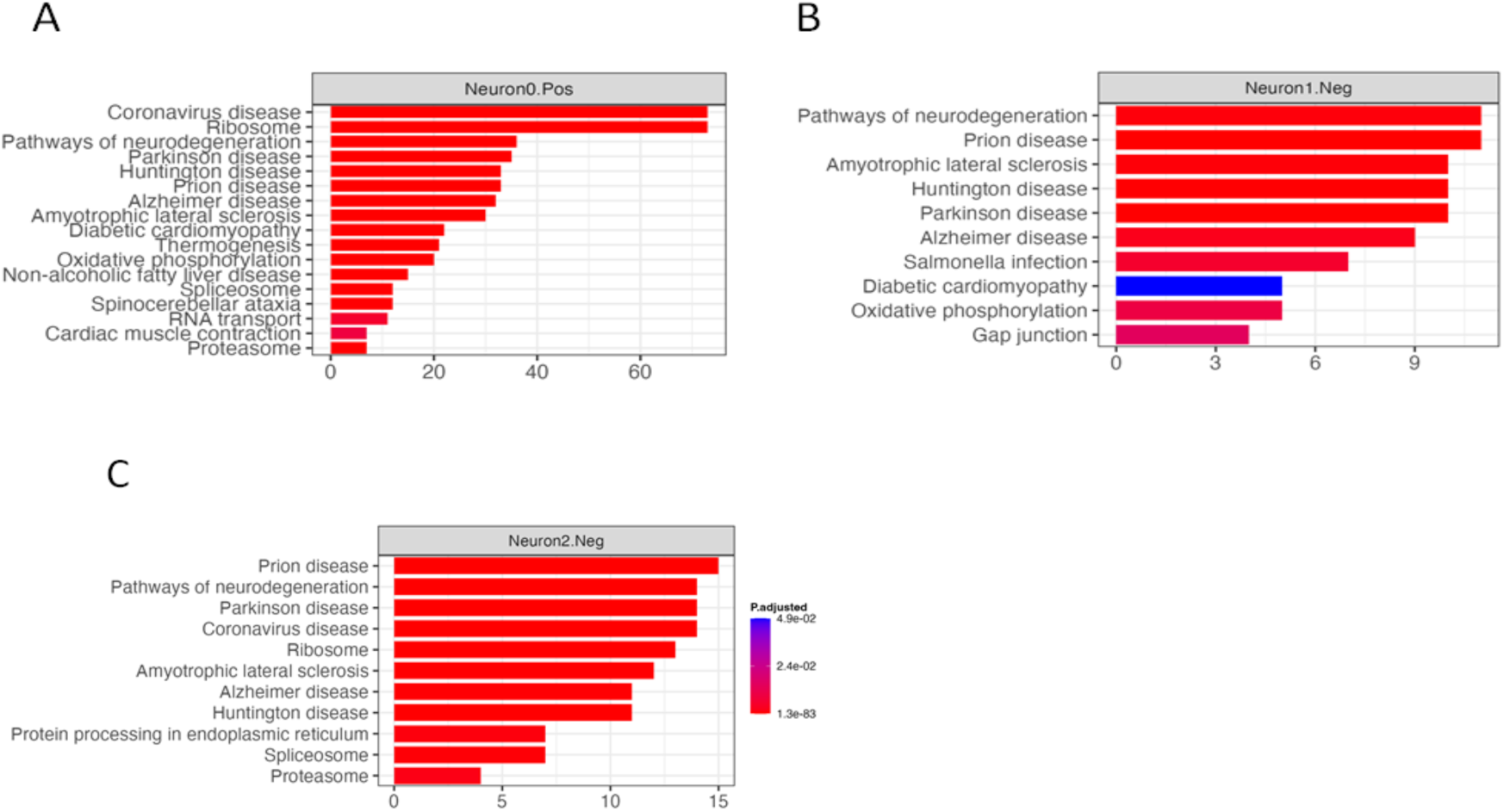
Neurodegenerative disease-relevant pathways were shown to be significantly altered in neuron subclusters 0, 1, and 2 of αSyn + PFF fed fish when compared to control fed siblings. Significant pathways upregulated in subcluster 0 included coronavirus, ribosome, neurodegeneration, Parkinson’s disease, Huntington disease, prion disease, Alzheimer’s disease and ALS, among others (A). Significant pathways downregulated in subcluster 1 include neurodegeneration, prion disease, ALS, Huntington’s disease, Parkinson’s disease, and Alzheimers disease, among others (B). Significant pathways downregulated in subcluster 2 included prion disease, neurodegeneration, Parkinson’s disease, coronavirus, ribosome, ALS, Alzheimer’s disease, and Huntington’s disease, among others (C).

## Discussion

The mistemplating and spread of αSyn pathology is a critical step in PD progression, but much of what surrounds this process remains opaque. This study describes the development of a zebrafish model of human αSyn neuronal expression that results in a slow and progressive degenerative phenotype. In young fish expressing αSyn, small changes in stimulated behavior are noted but with no detectable impact on overall survival or health. There has been previous work to establish a pan-neuronal human αSyn expressing zebrafish line which demonstrated that zebrafish can modify αSyn by phosphorylation and also that expression of αSyn protein caused no overt morphological deficit and did not impact survival in young fish^26^. These findings are consistent with our early studies, as is the finding that expression of the UAS: αSynT2AeGFP construct decreases with age.

Gene expression changes in these young fish expressing alpha-synuclein, as identified through single cell RNAseq, occur in neurons and changes are relatively minor. Microglia in these young fish show morphological features consistent with activation and phagocytosis. In addition, these young fish have a reduction in the number of TH-positive neurons in the telencephalon. As the fish age, those expressing αSyn develop progressive loss of HuC/D positive neurons in the gut as well as a loss of HuC/D protein in the brain, but this does not occur until after 6 months of age. It is significant that these fish appear relatively normal and have a slowly progressive phenotype that is highly age dependent because this mirrors what is seen in human disease caused by high expression of αSyn such as is the case in gene duplication^27,28^.

Because this transgenic line exhibits a very low level of early life toxicity, it serves as a unique model to study second hits that may initiate or perpetuate pathological spread and toxicity. To this end, we introduced a diet enriched with biologically active human αSyn PFF. Fish that both expressed αSyn and consumed PFFs for a period of 3 weeks had increased brain pSer129 labeling as well as fewer TH positive neurons and altered gene expression. These findings appear to recapitulate the initiation of pathological αSyn in the gut and spread to the brain as hypothesized in PD. This gut to the brain spread provides a novel method to determine the mechanisms of neuron to neuron spread and factors that alter it. This may provide insight into the reasons why PD progression is heterogenous and identify new targets for disease modification.

Support: *We thank the UCLA Institute for Quantitative and Computational Biosciences (QCB) for their support*

## Notes

**Competing Interests**: None

### Competing Interest Statement

The authors have declared no competing interest.

